# Electrostatics and Solvation: Essential Determinants of Chromatin Compaction

**DOI:** 10.1101/785634

**Authors:** A. Bendandi, S. Dante, S. Rehana Zia, A. Diaspro, W. Rocchia

## Abstract

Chromatin compaction is a process of fundamental importance in Biology, as it greatly influences cellular function and gene expression. The dynamics of compaction is determined by the interactions between DNA and histones, which are mainly mechanical and electrostatic. The high charge of DNA makes electrostatics extremely important for chromatin topology and dynamics. Besides their mechanical and steric role in the chromatin fibre, linker DNA length and linker histone presence and binding position also bear great electrostatic consequences. Electrostatics in chromatin is also indirectly linked to the DNA sequence: the presence of high-curvature AT-rich segments in DNA can cause conformational variations with electrostatic repercussions, attesting to the fact that the role of DNA is both structural and electrostatic. Electrostatics in this system has been analysed by extensively examining at the computational level the repercussions of varying ionic concentration, using all-atom, coarse-grained, and continuum models. There have been some tentative attempts to describe the force fields governing chromatin conformational changes and the energy landscapes of these transitions, but the intricacy of the system has hampered reaching a consensus. Chromatin compaction is a very complex issue, depending on many factors and spanning orders of magnitude in space and time in its dynamics. Therefore, comparison and complementation of theoretical models with experimental results is fundamental. Here, we present existing approaches to analyse electrostatics in chromatin and the different points of view from which this issue is treated. We pay particular attention to solvation, often overlooked in chromatin studies. We also present some numerical results on the solvation of nucleosome core particles. We discuss experimental techniques that have been combined with computational approaches and present some related experimental data such as the Z-potential of nucleosomes at varying ionic concentrations. Finally, we discuss how these observations support the importance of electrostatics and solvation in chromatin models.

**SIGNIFICANCE:** This work explores the determinants of chromatin compaction, focusing on the importance of electrostatic interactions and solvation. Chromatin compaction is an intrinsically multiscale issue, since processes concerning chromatin occur on a wide range of spatial and temporal scales. Since DNA is a highly charged macromolecule, electrostatic interactions are extremely significant for chromatin compaction, an effect examined in this work from many angles, such as the importance of ionic concentration and different ionic types, DNA-protein interactions, and solvation. Solvation is often overlooked in chromatin studies, especially in coarse-grained models, where the nucleosome core, the building block of the chromatin fibre, is represented as a rigid body, even though it has been observed that solvation influences chromatin even at the base-pair level.

## INTRODUCTION

Chromatin is formed through the winding of DNA around histone proteins. Its architecture allows the DNA to fit in the cell nucleus and supports structural integrity. Importantly, it helps the modulation of DNA compaction, allowing for efficient packing and unpacking mechanisms during transcription. The first level of packing is the nucleosome, composed of 147 DNA base pairs (bp), wrapped around the histone octamer (H2A, H2B, H3, and H4 dimers). The disordered domains of the histones – a total of ten histone tails – are particularly important for compaction, as they are highly charged and mobile. The other indisputably essential ingredient is the mechanical connection; for example, the length of linker DNA and the presence of high-curvature AT-rich segments (A-tracts) is known to influence internucleosomal interaction and impact on chromatin folding (1). Finally, the notion that the nucleosome is a solid object has been proven erroneous (2), indicating the importance of careful consideration of solvation effects in chromatin models. Our exploration of the role of electrostatics and solvation as determinants of chromatin compaction is paralleled with that of experimental techniques providing instrumental information for the validation and improvement of these models, focusing on methods that only minimally perturb the observed system.

In this work, we examine the fundamental importance of electrostatic interactions in chromatin, and their impact on fibre compaction and polymorphism. This brings us to an exploration of the, sometimes underrated, role of solvation in chromatin compaction modelling, and a presentation of quantitative results on the solvation of the nucleosome core particle, which support qualitative analyses found in the literature. Finally, we conclude with a discussion on experimental techniques that have been used in chromatin studies and present experimental measurements of electrostatic observables under varying ionic strength conditions.

## MATERIALS AND METHODS

### DLS and Zeta Potential measurements

Mononucleosomes assembled from recombinant human histones expressed in *E. coli* (two each of histones H2A, H2B, H3 and H4) wrapped by 147 base pairs of 601 positioning sequence DNA, were a product of EpiCypher (Durham, NC, USA). Mononucleosomes (100 μg total mass (protein and DNA), 54.6 μg protein) were delivered in 10 mM Tris-HCl pH 7.5, 1 mM EDTA, 25 mM NaCl, 2 mM DTT, and 20% glycerol. Aliquots (10 μg) were kept at the temperature of −20°C and defreezed before sample preparation. NaCl was a product of Sigma Aldrich (Saint Louis, MO, USA). Buffer solution containing NaCl in concentration 5, 10, 20, 50, 137, 250 and 1000 mM were prepared. For DLS and *ζ* measurement, 2.5 μg of nucleosomes were dissolved in 600 μL of buffer. Dynamic Light Scattering and potential measurements were collected with a NanoSizer Instrument (Malvern Panalytical, Malvern, UK). Three independent measurements for each sample were collected.

## ELECTROSTATIC INTERACTION IN CHROMATIN: AN INTRINSICALLY MULTISCALE PHENOMENON

Due to the high charge of the DNA, the electrostatic interaction rises as the de facto driving force of chromatin compaction. The charges present on the DNA backbone are partly neutralized by the winding of DNA around the histone core, especially through the effect of the histone tails, and partly through counter-ions present in the nuclear environment. The modelling of internucleosomal interactions using reductionist analytical potentials, which omit the explicit role of histone tails, can cause secondary, but still relevant, electrostatic effects to be overlooked. Considering the biological importance of different ionic types, Mg^2+^ is particularly significant, as it has been found to promote nucleosome condensation and aggregation and could promote linker DNA bending, because in its presence interactions of first and third neighbouring nucleosomes are boosted (3). Tetravalent cations on the other hand require lower concentrations to induce compaction. In (4), systems of 1-10 nucleosome core particles (NCPs) were studied using a coarse-grained model in order to study the effects of monovalent, divalent, and trivalent cations on these structures. It was observed that an increase in K^+^ ions amplified the repulsive internucleosomal electrostatic interaction; increasing Mg^2+^ concentration caused partial aggregation, and an increase in COHex^3+^ ions triggered a strong mutual internucleosomal attraction in 10 NCP systems, therefore showing that the aggregation of NCPs is different under the effect of different types and concentrations of counterions.

Multivalent ions and the effect of their distribution around NCPs on chromatin conformation have been studied also in (5), using a mean-field PBE approach, with an emphasis on shielding charges, which aggregate particularly around DNA and the exposed parts of the histone tails. The fact that a surface needs to be exposed to solvent in order for ions to bind on it makes ion-caused electrostatic screening and ion-chromatin interactions in general directly dependent on compaction. Calculations showed that the enhanced screening due to divalent ions might not only be because of their higher charge, but also because they form a denser layer of counterions around the NCP and fluctuations in this layer are correlated to different fibre conformations. This makes even more evident the fact that the topology of compaction is a key determinant for chromatin-ion interaction. It was observed in these simulations that the shielding charge arising from both monovalent and divalent ions was linearly correlated with the ionic strength of the solution.

In the study of structures as large and complex as chromatin, it has been proposed in (6) that implicit solvent Generalized Born (GB) simulations would be preferable to traditional fully explicit molecular dynamics (MD), in order to circumvent computational limitations. However, standard GB scales poorly with the number of solute atoms and, in this work, a multiscale atomistic GB model that incorporates improvements in the electrostatic calculations is presented, the accuracy of which was evaluated through point-by-point comparison with Poisson Boltzmann Equation (PBE) calculations. Taking advantage of the natural hierarchical organization and charge distribution of chromatin, they used approximate point charges to calculate electrostatic interactions between distant points in a 40-nucleosome structure, containing approximately 1 million atoms, focusing particularly on the behaviour of the histone tails. This approach proved the existence of viable alternatives that drastically reduce the cost of conformational sampling in very large structures.

Linker DNA length is extremely important for chromatin compaction, not only for mechanical but also for electrostatic reasons. Determining how linker DNA influences chromatin topology, and how its length and sequence can affect compaction has been the subject of many studies and speculations. In the work of (1), for instance, the presence of so-called A-tracts, DNA fibres where multiple A-T pairs are present in a row, and its influence on DNA rigidity, and therefore on chromatin fibre flexibility, are examined. It has been observed that the presence of A-tracts causes DNA bending angles of up to 90°, and that these particular segments are often found in linker DNA (7). The direction of bending of the linker DNA is also relevant for compaction: for example, when DNA bends inwards at the exit sites from the NCP the resulting structures are more compact compared to the opposite case and give rise to zig-zag configurations and closer overall nucleosome proximity. It is evident that linker DNA length is certainly of great importance when it comes to chromatin topology; however, its role is not immediate, in that the really important parameters for packing are the DNA binding angles, which are influenced by linker DNA length through topological and persistence length constraints.

The presence of the linker histone H1 (or H5 in avian chromatin) is also a key for compaction. This histone is not always present in nucleosomes, and its position can vary on or off the nucleosome dyad axis (8). The H1/H5 changes the orientation and flexibility of linker DNA, making contacts with both entering and exiting strands. When two or more nucleosomes in sequence are bound to H1 histones, rigid structures termed DNA stems are formed, which present straighter linker DNA and reduced separation angle between the entering and exiting DNA; the latter effect is more pronounced in chromatin configurations with long linkers (9). The increased rigidity of DNA because of the formation of DNA stems is mitigated by the dynamic nature of H1/H5 binding and unbinding on nucleosomes (10).

One could not conclude a discourse on chromatin electrostatics without mentioning the effect of the histone tails, which have been found to promote stability of the linker histone on the NCP. In some models, histone tails are modelled as a series of beads with one positive charge per bead (4, 5, 11). It was seen by (12) that certain histone tail configurations promote DNA bulging at entry and exit sites, possibly contributing to the formation of twist defects in the nucleosomal DNA, which are important, among other reasons, because their presence causes the formation of nucleosomes with 146 bp instead of the usual 147 (13), due to overwinding and stretching of the DNA (14). They also speculated that the presence of arginines and lysines might impose constraints on histone tail motion because of attractive electrostatic interactions. Contacts between DNA and histones were seen to be dominated by the histone tails, making up 60% of protein-DNA interactions in the nucleosome, rapidly wrapping around the DNA (in (12), it was observed that they do so in the first 20ns of the simulation). In another study, the N-terminal of the H4 histone tail was observed to interact with the “acidic patch” present on the surface of adjacent nucleosomes, a small groove formed by eight residues, six belonging to H2A and the remaining to H2B, which constitutes a region of highly negative charge density on the nucleosome surface, serving as a hot-spot for DNA-binding proteins and histone tails (15–17). Throughout 1 μs-long MD simulations, the NCP is seen to be very stable in dynamics, in contrast to histone tails and linker DNA: large scale unwrapping or opening of NCP DNA were not observed, even when simulations were performed in 1M salt concentration, under which conditions they are known to occur (18). This indicates that such phenomena might take place on longer time scales. Of particular interest are the histone H3 tails, which have been suggested by experiments (19) to form stable folded structures, and even to potentially compete with other DNA-binding proteins, affecting accessibility of epigenetically modified sites in the minor grooves.

It has already been mentioned that the presence of A-tracts can change the curvature of DNA, causing the minor grooves to be narrower than those in segments with lower curvature, and locally enhancing negative electrostatic potentials (20). In (20), PBE calculations were performed on DNA, showing that the electrostatic potential caused by the DNA backbone had intensity peaks inside the major and minor grooves. The position of these peaks correlates with the positions of arginine residues on the histone core. Previously observed binding preference for arginines over lysines in minor grooves, and especially in the narrower ones, was partly explained via a combination of electrostatic and desolvation effects.

## THE ROLE OF SOLVATION IN CHROMATIN COMPACTION

The role of the solvent in biomolecular interactions is known to be crucial. In part, this is because of solvent-mediated electrostatic effects, that is the screening of the water molecules and that of the ions in solution. In addition, there is the so-called cavity formation phenomenon, which penalizes the occurrence of solvent-excluded regions. Chromatin spatial arrangement, due to NCP charge, size and porosity, is expected to be particularly affected by these phenomena, which must be accurately considered. It has already been described that the formation of the fundamental unit of chromatin, the nucleosome, is carried out by the complexation of the negatively-charged DNA polymer with the positively-charged histone protein octamer. If investigated at the molecular level, this process is governed by a number of interactions such as hydrogen-bonds, salt-bridges, and water-mediated interactions occurring along the positively-charged arginine anchors that intercalate deep inside the minor grooves of DNA facing the histone core (16, 21). When it comes to histone core-DNA electrostatic interactions, it is known that (22) every nucleosome presents 14 non-covalent histone-DNA contacts, at the sites of arginine residues.

Solvent exposure affects electrostatic interactions at the nucleosome level: compared to H3 and H4 histones, the two H2 variants are more solvent exposed, making them more accessible to chromatin-binding proteins as well (6). Specific ion binding sites and their location on the nucleosome are also of particular interest, and they can be studied using electron density maps in combination with chemical information (14). It has been observed that sodium preferentially condenses around regions rich in solvent accessible acidic residues, especially in areas with two or more acidic residues in close proximity (2). It is also speculated that, in chromatin fibres exhibiting high compaction, internucleosomal electrostatic repulsion could be reduced in intensity because of an increased neutralization of the DNA backbone charge by the neighbouring histone cores and counterion screening.

The idea that the nucleosome is an impermeable object has been proved erroneous (2); in this work, it was seen that mobile ions are able to reach the NCP inner core because of high levels of local solvation (more than 1000 water molecules). This led to the conclusion that the local value of dielectric constant in the region facing the histone core is larger than expected. The authors also looked into the mobility of water molecules on the first hydration layer of the nucleosome and, as expected, found them to be less mobile than bulk water molecules. Through detailed visualization of structured water at the protein-DNA interface, they also found that water molecules not only contribute significantly to the stability of DNA binding but also adapt histone surfaces to conformational variations of DNA, facilitating nucleosome dynamics. All-atom electrostatics calculations were conducted (2) and compared to PBE calculations, observing a slight inconsistency between the two. PBE predicts that the most significant contribution to DNA charge neutralization comes from the enhancement of the electric field and that it is a result of the tight wrapping of the DNA around the histone core. Their results indicate that close condensation of ions around the nucleosome can significantly reduce the short range effect of the nucleosomal charge, having as a natural consequence the facilitation of chromatin close packing.

In another work concerning NCP solvation (14), the solvent-accessible surface area (SASA) of nucleosome crystals with 147 bp and 146 bp was investigated. NCPs with 147 bp were found to possess a SASA of approximately 74 Å^2^, which is distributed mostly in the cavities within the histone octamer and in the space between it and the DNA. The primary hydration layer of the NCP was found to contain slightly more than 2000 water molecules, the positions of which were found to largely correspond to the positions of A-tracts, especially in the vicinity of the minor groove. In this work, water has been shown to be important in the two main mechanisms of protein-DNA recognition: direct readout (nucleotide chemically specific bonds) and indirect readout (sequence-dependent conformational features of DNA recognized by sterically complementary protein contacts). Structures termed “spines of hydration” were also observed, in which water molecules bind regularly to adenine N3 and thymine O2 atoms (23). Structural analyses have shown that the phosphate groups are the most strongly solvated components of the DNA (24, 25).

To analyse the porosity of the nucleosome, particularly described in (2), we have conducted a quantitative study on the nucleosome crystal structure (PDB code 1kx5 (14)) using NanoShaper (26) interfaced with VMD (27), providing the values of the Surface to Volume Ratio (SVR), the number of cavities and pockets. We observe an SVR of 0.387 Å^−1^, which reflects a quite high porosity(28), 11 closed cavities with volumes ranging from 20.62Å^3^ to 55.75Å^3^ and 31 pockets. We visualize the channel traversing the nucleosome core, which significantly impacts on NCP accessibility to water and ions. Our results are consistent with previous qualitative analyses mentioned in this section, and indeed indicate that the nucleosome is highly solvated and porous. We have also constructed an electrostatic map of the nucleosome, using data from DelPhi on the potential and constructing the SASA of the nucleosome with NanoShaper (videos of the full 3D structure found in Supplementary Material), as seen in Fig.1, where it is possible to clearly see, among other features, the position of the acidic patch (residues E56, E61, E64, D90, E91, E92 of H2A and E102, E110 on histone H2B (15)), and the highly charged histone tails, both key elements in chromatin compaction and chromatin interaction with DNA-binding proteins. We also observe a minor acidic region, on the surface of histone H4.

**Figure 1:**
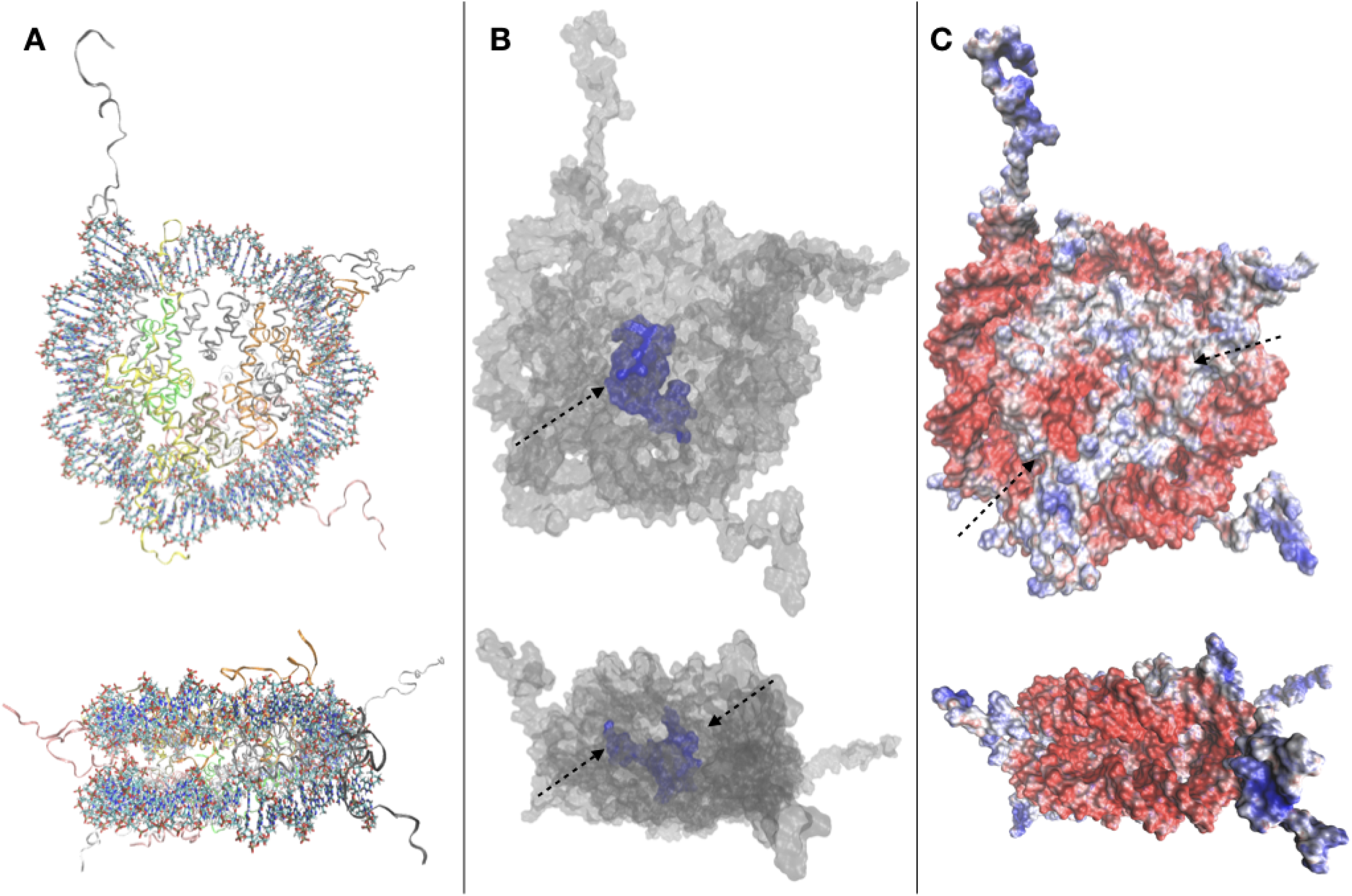
**A.** Top and side view of the 1kx5 crystal structure. **B.** Top and side view of the SES of 1kx5, constructed with NanoShaper (26) and visualized via VMD. The channel traversing the histone core is represented in blue together with an adjacent open cavity and is indicated by an arrow. On the side view, the entrance and exit of the channel can be seen, indicated by arrows. **C.** Electrostatic map of the SES of 1kx5. Areas of negative surface potential are indicated in red and areas of positive surface potential in blue. The acidic patch is indicated by an arrow on the histone core. Another minor acidic region, composed by fewer residues on the surface of histone H4, is also highlighted. Most of the exposed regions of the histone core are electrically neutral, with the acidic patch representing the main exception. Remaining positive charges of the histone core are buried, due to the binding of encircling DNA. We also note positive charges on the histone tails, and strong negative charges on the DNA backbone.

Non-invasive experimental observations of quantities related to electrostatics and solvation in chromatin can be done via DLS and Zeta Potential measurements. They can be done on single nucleosomes and under varying ionic conditions. To our knowledge, this is the first application of this technique on nucleosomes. Fig.2 summarises the results, displaying both size d and *ζ* measured at different salt concentrations. In the interval between 5 and 250 mM NaCl concentration, the absolute value of *ζ* was found to decrease monotonically from 45±7 mV to 17±7mV when the salt concentration increases. This result is in good agreement with the *ζ* values extrapolated from electrophoretic mobility and reported in the literature (29). In the same NaCl concentration range, the size measured by DLS varied between 1.9±1.3 and 4.6±1.7 nm. At the highest concentration (i.e., 1 M NaCl) large aggregates were present in the sample (d=109±23 nm), and the corresponding *ζ* was −4.6 mV±1.8. As we mentioned before in this work, in such high concentrations nucleosomes have been observed to be unstable, and disassembly is possible, explaining the formation of such aggregates. Interestingly, Zeta Potential measurements can be compared to the average, taken at a suitable distance from the Solvent Excluded Eurface (SES) of the solute, of the electrostatic potential obtained, for instance, by solving the PBE. This can be done by exploiting already existing features of PBE solvers, such as DelPhi (30).

**Figure 2.**
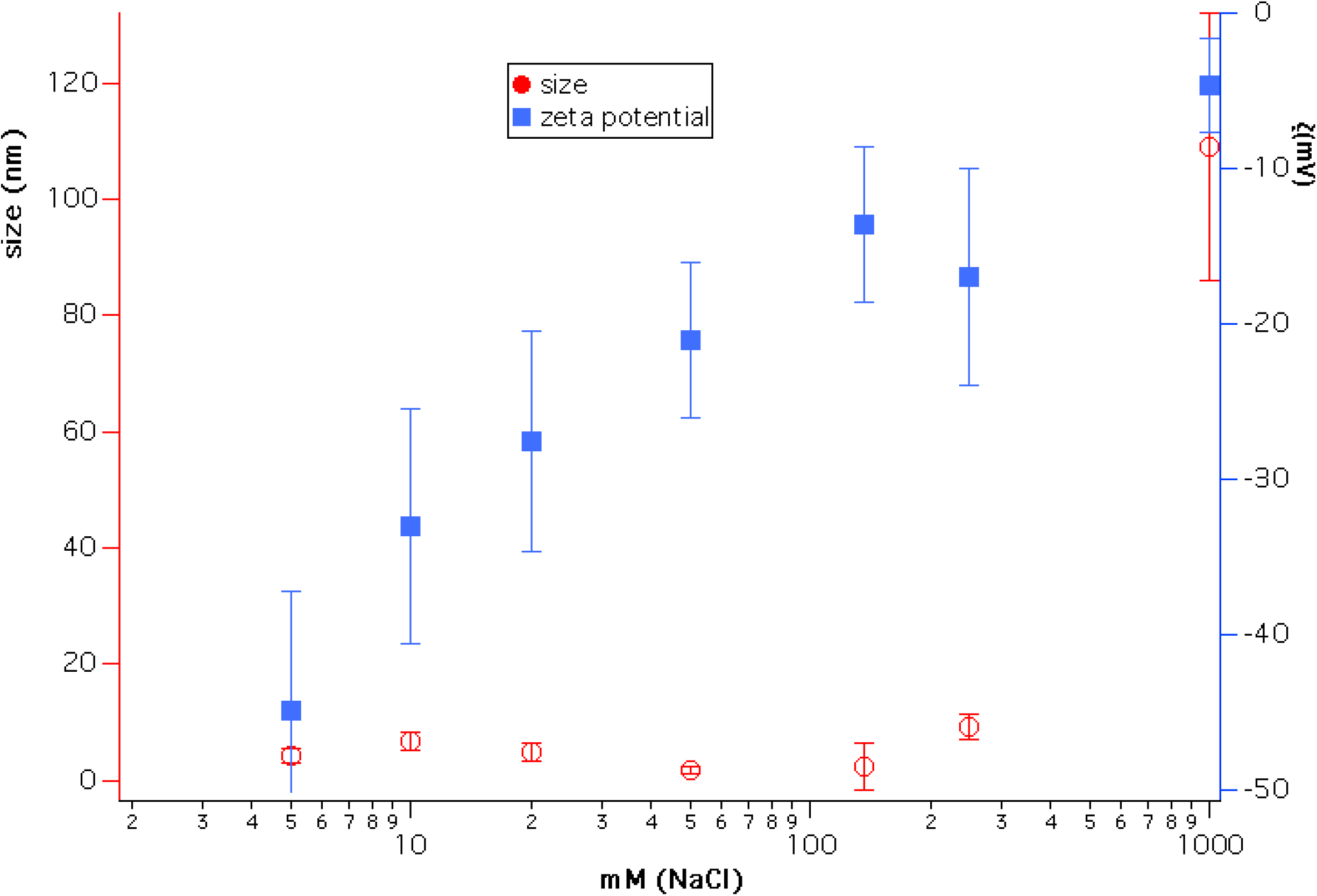
Zeta potential and particle size measured under varying ionic conditions. A clear trend is seen in the decrease of *ζ* as a function of NaCl concentration. At high concentration (1M), aggregates are observed in the sample, possibly owing to the disassembly of NCPs.

## EXPERIMENTAL STUDIES OF CHROMATIN: FROM THE NUCLEOSOME TO THE NUCLEUS

We have already highlighted here the multiscale nature of chromatin, and explored the main determinants of its compaction. Given its degree of complexity, it is evident that simulations alone would be insufficient for a thorough study of chromatin. Indeed, the interplay between simulations and experiments is crucial in this field and has given rise to breakthroughs that would have been impossible without the combination of the two approaches. Experimental investigations of chromatin can be carried out at different scales, similarly to computational approaches. Having already mentioned some experimental results validating computational models, we specifically look into some of the experimental techniques used, in both small and large scales, from the NCP up to entire nucleus.

Starting from the building block of chromatin, the nucleosome, experiments have been carried out to determine its crystal structure, with continuing endeavours starting from (31), in which a 2.8Å resolution structure of the NCP was obtained via X-ray crystallography, using reconstituted nucleosomes. In Widom’s work, many of the structural elements of the nucleosome were uncovered, such as the the number of base pairs wrapped around the octamer, which were unknown despite the octamer histone structure had already been observed. The histone tails and their structural role have also been studied to great extent in (31). Since then, further structures with 147 bp (14) and 146 bp (32) have been observed. The study of sub-structures such as the histone tails and of site-specific interactions (33) in more detail, required the use of NMR (14). In latest years, there has been growing interest on the study of NCPs using Cryo-EM. The sample preparation protocols involved in this technique make it faster to use for structural studies than X-ray crystallography. Cryo-EM provided information on custom-made NCPs in studies relevant to DNA binding protein-NCP interactions (34) and also on interactions of the NCP with components of the nuclear environment, such as the nuclear pore complex (35). The orientation of NCPs has also been observed by Cryo-EM in a recent study, where it is stated that in the most common arrangement of a pair of NCPs they are placed in parallel, facing histone octamers (36).

X-ray crystallography provides structures with atomic resolution, which are key for atomic-level studies. However, this approach has some limitations; it fails to provide good information on the more mobile domains of the NCP, and it cannot be used for large oligonucleosomes (the largest structures that have been crystallized to date are tetranucleosomes (37, 38)). In order to circumvent these constraints, one can turn to scattering techniques. Small Angle X-ray Scattering (SAXS) studies have looked into the issue of whether the histone tails protrude into the solvent surrounding the NCP or associate with DNA at physiological salt conditions. The histone tails are notoriously hard to resolve in crystallography because of their size and intrinsically mobile nature, and structures including them usually come from NMR measurements (19, 39, 40). Using SAXS however, it is possible to indirectly observe whether the histone tails are solvated or adherent to the DNA, by measuring changes in the overall structure size. (41) have applied SAXS to study histone tails as well, focusing on the structural details of internucleosomal interactions and the effects that histone tails have on them. Often SAXS has been used in conjunction to other techniques to correlate structural to dynamical data. In (42) SAXS, Forster Resonance Energy Transfer (FRET), and MD were used to dissect the sequence-dependent DNA unwrapping mechanism. Fluorescence Correlation Spectroscopy (FCS) has been used in a work by (4) to estimate NCP stacking energy. In this combined experimental and theoretical work, model parameters were tuned based on comparison with single molecule FCS and SAXS data, which also showed that histone tails facilitate NCP stacking by acting as bridges between NCP surfaces. FCS data was also used by (43) to tune the parameters of an MC model of nucleosome arrays under the influence of external forces.

Moving on from NCPs to larger structures, nucleosome arrays are the next step; besides SAXS (44), Atomic Force Microscopy (AFM) in liquid has also been used to study arrays of varying lengths. The advantage of using this technique for chromatin is twofold: there is the possibility of taking many measurements, making it good for statistical purposes; and it allows for the study of electrostatic and related interactions, such as differences in ionic concentration. The importance of ionic interactions with chromatin has naturally gained the attention of the experimental community. Experimental studies (5) have shown that Mn^2+^ ions bind to the major DNA groove near CG pairs. In (45) AFM was used to measure the changes in chromatin topological conformations depending on salt levels in the environment. Studying NCP arrays in varying salt concentration revealed that array compaction has a non-monotonic salt dependence. Increasing salt concentration induces partial screening of the charges of the DNA backbone, therefore reducing the electrostatic interactions between DNA and histones, directly impacting on compaction. The stability of mononucleosomes has also been investigated in correlation with salt concentration (46): in low to intermediate salt regimes they observed some partially disassembled states (as also studied computationally in (47)) where H2A/H2B histone dimers partially dissociate from the NCP. Regarding the mechanical properties of chromatin, DNA stiffness was observed to be salt-dependent as well, in accordance with other experimental and computational studies (13, 20, 48, 49); the persistence length was seen to increase at higher ionic concentrations.

Optical microscopy, a field traditionally tied to biological applications, is a natural candidate for chromatin studies, due to the advances in resolution obtained by super-resolution techniques, and to the fact that label-free optical microscopy methods have been on the rise for the past decade. Experiments using the single molecule super-resolution microscopy technique STORM (50) have observed units of chromatin organization termed by the authors *clutches*, heterogeneous groups of various sizes. The size of the clutches has been speculated by Ricci et al. to be related to the pluripotency capacity of each cell, and the median number and nucleosome density in the nucleus was found to be cell-specific. From longer nucleosome arrays to chromatin fibre, other super-resolution techniques, such as Photoactivated Localization Microscopy (PALM) have been used to extrapolate chromatin topology in the nucleus from nucleosome dynamics. Label-free techniques are also used to study chromatin at the nuclear level, such as Circular Intensity Differential Scanning (CIDS) (51). SAXS and Cryo-EM have also been used in structural analysis of the fibre up to the chromosome level (52–55). Experimental validation has been attempted also for some among the most exotic theoretical models proposed for chromatin, namely those hypothesising fractal globules. Fractal globules have been observed experimentally in Hi-C experiments (56) and Small-Angle Neutron Scattering (SANS) experiments (57, 58). SANS has been considered a good technique for experiments looking for fractal structures in the nucleus because of its extended spatial range, from approximately 15 nm to 10 μm (59). The important question tackled by works on this topic is the way in which fractal states with stable long-lived properties are formed.

## CONCLUSION

Electrostatics in chromatin is involved in an intricate combination of different mechanisms and its role in compaction and chromatin remodelling is of paramount importance. The highly negative charge of DNA is partially neutralized by its direct interaction with histones (including the effects of histone tails and the linker histone), but fibre electrostatic stabilisation is achieved through a combination of this effect with long-range electrostatics and ionic screening. Simulations in which ionic interactions with chromatin at the NCP level are treated more accurately would be a great improvement to existing approaches. In addition, based on the results of (2), we conclude that a more accurate representation of the nucleosome core is crucial when performing these analyses, since solvation has proved to be a very important factor in nucleosome behaviour, and therefore in chromatin compaction, whereas neglecting these effects would hamper a correct understanding of chromatin compaction. We support this claim with some numerical data on the structure and solvation of the NCP, which show that it is highly porous and solvated, and with zeta potential measurements at different ionic strengths. In summary, we have presented on the role of electrostatics and solvation as the driving mechanisms of chromatin conformational changes and equilibria, and showed how modern physics-based models of chromatin compaction must account for these phenomena. To complement this overview, we also presented some representative experimental approaches to study chromatin structure and dynamics, at both small and large scales.

## AUTHOR CONTRIBUTIONS

AB performed most of the reviewing and writing tasks. SD and AD organised and discussed the experimental part. SD performed the experiments. SRZ worked on the analysis of the role of solvation. WR and AB decided the organisation of the manuscript and checked the consistency of the work. All Authors reviewed and checked the manuscript.

